# Sympatric and allopatric differentiation delineates population structure in free-living terrestrial bacteria

**DOI:** 10.1101/644468

**Authors:** Alexander B. Chase, Philip Arevalo, Eoin L. Brodie, Martin F. Polz, Ulas Karaoz, Jennifer B.H. Martiny

**Affiliations:** Department of Ecology and Evolutionary Biology, University of California, Irvine, CA, USA; Earth and Environmental Sciences, Lawrence Berkeley National Laboratory, Berkeley, CA, USA; Department of Civil and Environmental Engineering, Massachusetts Institute of Technology, Cambridge, MA, USA; Department of Environmental Science, Policy, and Management, University of California, Berkeley, CA, USA; Center for Marine Biotechnology and Biomedicine, Scripps Institution of Oceanography, University of California, San Diego, CA, USA; Department of Ecology and Evolutionary Biology, University of Chicago, Chicago, IL, USA

## Abstract

In free-living bacteria and archaea, the equivalent of the biological species concept does not exist, creating several barriers to the study of the processes contributing to microbial diversification. As such, microorganisms are often operationally defined using conserved marker genes (i.e., 16S rRNA gene) or whole-genome measurements (i.e., ANI) to interpret intra-specific processes. However, as in eukaryotes, investigations into microbial populations must consider the potential for interacting genotypes among individuals that are subjected to similar environmental selective pressures. Therefore, we isolated 26 strains within a single bacterial ecotype (equivalent to a eukaryotic species definition) from a common habitat (leaf litter) across a regional climate gradient and asked whether the genetic diversity in a free-living soil bacterium (*Curtobacterium*) was consistent with patterns of allopatric or sympatric differentiation. By examining patterns of gene flow, our results indicate that microbial populations are delineated by gene flow discontinuities and exhibit evidence for population-specific adaptation. We conclude that the genetic structure within this bacterium is due to both adaptation within localized microenvironments (isolation-by-environment) as well as dispersal limitation between geographic locations (isolation-by-distance).

## INTRODUCTION

In eukaryotes, populations are typically defined as groups of interbreeding individuals within a species residing in the same geographic area (Mayr, 2001). Geographically-distinct (i.e., allopatric) populations are also often genetically distinct because of reduced gene flow, or the exchange of genetic variation, between populations of the same species. However, in microorganisms, the equivalent of the biological species concept does not exist, creating several barriers to the study of the fine-scale genetic structure of microbial populations and thus, the processes contributing to microbial diversification (Chase and Martiny, 2018; Shapiro *et al.*, 2016; Rocha, 2018).

The first of these barriers is that the genetic resolution delineating a microbial population is unclear. In eukaryotes, populations are, by definition, genetic units belonging to the same species, but defining a prokaryotic species remains challenging (Ward *et al.*, 2008). Nonetheless, there is evidence for geographically-distinct, genetically-diverged groups of bacteria and archaea. Several studies have shown that the genetic similarity of closely-related microbial individuals are negatively correlated with geographic distance across continental and global scales (Andam *et al.*, 2016; Choudoir *et al.*, 2016; Whitaker *et al.*, 2003; Zwirglmaier *et al.*, 2008). This pattern is consistent with isolation-by-distance, whereby dispersal limitation contributes to reproductive isolation over geographic distances (Wright, 1943). Further, in some cases, these geographically-localized genetic clades appear to be adapted to local environmental conditions, as individuals within these clades can differ in their temperature (Choudoir and Buckley, 2018), nutrient (Johnson *et al.*, 2006), or habitat preference (VanInsberghe *et al.*, 2015). However, the degree of divergence between genetic clades in such studies is usually quite high (<90% genome-wide average nucleotide identity), indicating they may not represent intra-species relationships (Jain *et al.*, 2017). These genetic units would seem to be much broader than populations, or groups of individuals with the potential for contemporary interactions and exchange of genetic material (Cordero and Polz, 2014). Therefore, a focus on much more closely-related microorganisms is needed to investigate the processes responsible for initial diversification.

A second, related obstacle is recovering genetically-similar individuals of the same species, however defined. Population genetic studies of eukaryotes typically characterize the genetic diversity among many individuals from a variety of geographic locations. For microbes, this sampling design requires reliable isolation of closely-related strains (but see (Kashtan *et al.*, 2014)), which can be difficult in highly diverse microbial communities such as soil. Finally, even if a sample of closely-related individuals can be collected, a third barrier is quantifying the exchange of genetic variation (i.e., gene flow) between individuals. For prokaryotes, the exchange of genetic material is mediated through genetic recombination, whether homologous recombination or horizontal transfer of entirely new genes. However, the asexual nature of prokaryotes makes it a challenge to quantify this process, particularly among closely-related individuals. The more closely-related two genomes are, the more difficult it is to distinguish between differences caused by vertical inheritance and recombination (Ravenhall *et al.*, 2015).

In aquatic (Cui *et al.*, 2015) and host-associated (Sheppard *et al.*, 2008) systems, many of these obstacles have been addressed. In these environments, geographic proximity does not appear to be the most important factor in structuring microbial populations as typically observed in plants and animals. Indeed, an increasing number of studies find several, distinct genetic clades co-occurring in the same geographic location (Hunt *et al.*, 2008; Cohan, 2001; Chase *et al.*, 2017; Whitaker *et al.*, 2005). For instance, the thermophilic archaeon, *Sulfolobus*, exhibited strong barriers to recombination between sympatric clades within a hotspring (Cadillo-Quiroz *et al.*, 2012). Such evidence suggests that the genetic structure of microbial populations is influenced less by divergence among geographically-distinct (allopatric) groups, and more by ecological differentiation (isolation-by-environment (Wang and Bradburd, 2014)) among co-occurring (sympatric) groups (Polz *et al.*, 2013). Thus, we might need to abandon the idea of defining microbial populations *a priori* based on geography (as done for larger organisms) and, instead, focus first on the emerging genetic structure among closely-related individuals (Arevalo *et al.*, 2019).

Soils are highly heterogeneous systems where differences in microhabitats can contribute to environmental variation over many spatial scales (Ranjard and Richaume, 2001; Nannipieri *et al.*, 2003). For this reason, one might expect that allopatric differentiation might be more evident for soil bacteria than that observed in aquatic environments. Indeed, some soil fungi exhibit strong population structure at regional spatial scales (Amend *et al.*, 2010; Branco *et al.*, 2015). Therefore, we asked whether population structure in a free-living soil bacterium was consistent with patterns of allopatric or sympatric speciation. To do so, we investigated the abundant leaf litter taxon, *Curtobacterium* (Chase *et al.*, 2016), which is relatively easy to culture from the leaf litter layer of soil. Previously, we demonstrated that *Curtobacterium* encompasses multiple ecotypes, or fine-scale genetic clades that correspond to ecologically relevant phenotypes (Chase *et al.*, 2018). Here, we concentrated on the genetic diversity within a single ecotype, *Curtobacterium* Subclade IB/C, a unit that might be considered equivalent to a species designation (Chase *et al.*, 2018). Specifically, we examined 26 strains (with identical full-length 16S rRNA regions and ≥97% genome-wide average amino acid identity) from a regional climate gradient, along with two closely-related strains isolated across continental distances. We hypothesized that soil bacteria would exhibit a pattern intermediate to that of aquatic free-living bacteria and archaea and soil fungi. In particular, we expected that sympatric populations of soil bacteria may exist within a particular geographic location, while also exhibiting a pattern consistent with allopatric differentiation among locations. Such a pattern would indicate that the genetic structure within this bacterium is due to both adaptation within localized microenvironments (isolation-by-environment) as well as dispersal limitation between geographic locations (isolation-by-distance).

## RESULTS

### Evolutionary History within a *Curtobacterium* ecotype

We identified 26 strains from a *Curtobacterium* ecotype, subclade IB/C, that share ecologically-relevant genotypic and phenotypic characteristics. These traits include the ability to degrade polymeric carbohydrates (i.e., cellulose and xylan), the degree of biofilm formation, and temperature preference for both growth and carbon degradation (Chase *et al.*, 2018). These strains were previously isolated from leaf litter, the top layer of soil, at four geographic locations from a regional climate gradient in southern California (Supplementary Table 1). All analyzed strains have identical full-length 16S rRNA regions and share high sequence identity with ≥94.6% genome average nucleotide identity (ANI) and ≥95.3% genome average amino acid identity (AAI), congruent with previous observations for defining discrete sequence clusters within natural microbial communities (Rodriguez-R and Konstantinidis, 2014). We also included two additional strains from subclade IB/C that were isolated from leaf litter in Boston, MA to provide varying geographic scales (ANI_[MEAN SIMILARITY]_ = 94.9%; AAI_[MEAN SIMILARITY]_ = 95.8%).

To examine whether genetically-similar strains within the IB/C subclade clustered by geographic location, we reconstructed the phylogenetic relationship among the strains using the core genome (Fig. 1A). The core genome phylogeny revealed highly structured genetic lineages; however, clusters contained strains isolated from a variety of geographic locations. While one strain from Boston, MA formed the outgroup, the other Boston strain was highly similar to a grassland strain from Loma Ridge, CA. At the regional scale within the climate gradient, most of the grassland strains clustered together, while strains from the scrubland and Salton Sea leaf litter communities were dispersed throughout the tree.

**FIG 1.**
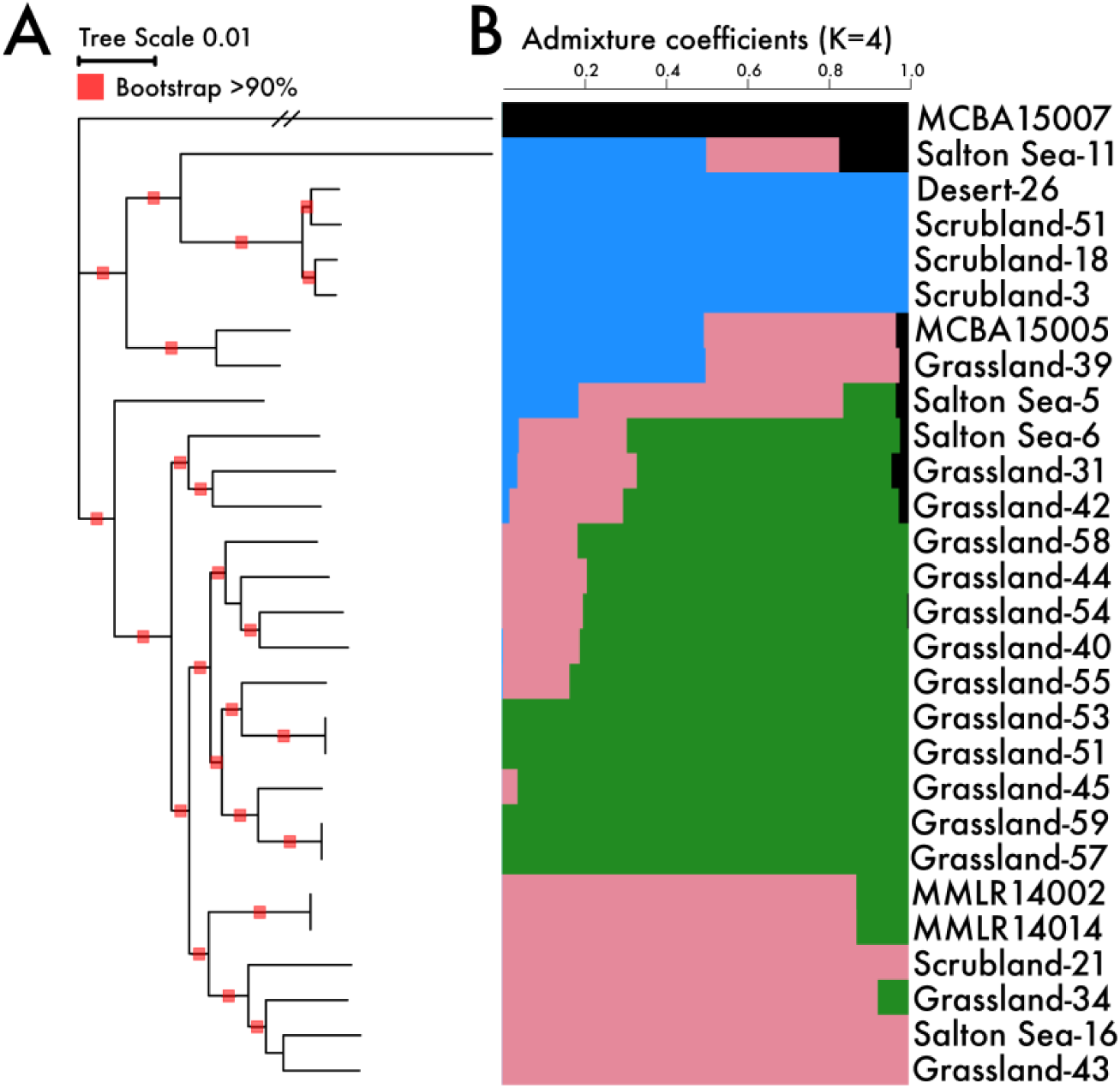
(A) Phylogeny of the *Curtobacterium* ecotype, subclade IB/C, from a core genome alignment. (B) Ancestral population structure estimated from admixture analysis. Bar plots reflect the proportion of an individual genome that originate from estimated ancestral gene pools (K = 4). Genome names designate the site of isolation along the climate gradient except for MCBA = Boston and MMLR = Grassland isolate from 2010.

Phylogenetic analyses alone cannot delineate population structure as it is necessary to account for both vertical descent and contributions from shared ancestral gene pools. Therefore, we supplemented the phylogenetic analysis by computing ancestry coefficients for each strain across the core genome using a STRUCTURE-like (Frichot and François, 2015) analysis (Fig. 1B). The most probable number of ancestral gene pools (K=4) contributing to the proportion of an individual genome (see Materials and Methods) demonstrated high congruence with the phylogenetic analysis. For example, an outgroup strain originating from Boston, MA exhibited little evidence for mixing with most of the climate gradient strains in CA across continental scales (Fig. 1B). Within the regional climate gradient, we detected three ancestral gene pools that may represent finer population structure across ecologically-similar strains in ecotype IB/C.

### Gene Flow Delineates Bacterial Populations

Although STRUCTURE-like analyses can provide insights into the genetic structure among divergent lineages, populations (defined as groups with the potential to exchange genetic material) must be resolved by examining patterns of gene flow. However, in asexual organisms, measurements of homologous recombination can be overestimated when individuals are closely related as distinguishing between recombination and point mutations is difficult (Ravenhall *et al.*, 2015). Further, other forms of horizontal gene transfer can be ecologically relevant as well (van Elsas and Bailey, 2002). To address these limitations, we employed a novel method, PopCOGenT, that attempts to detect all recent recombination events between pairs of strains (Arevalo *et al.*, 2019).

To distinguish between vertical descent and homologous recombination in structuring populations, we used PopCOGenT to estimate the degree of recombination among the genomes. This analysis revealed three recombining populations that are evident as highly isolated clusters in the network (Fig. 2). One of the populations (population 2) was restricted to a single location (in the grassland site). The other two populations included strains from multiple sites along the climate gradient; for example, population 3 contained strains isolated from the grassland, scrubland, and Salton Sea leaf litter communities, which are geographically separated by 177 km (Supplementary Table 2).

**FIG 2.**
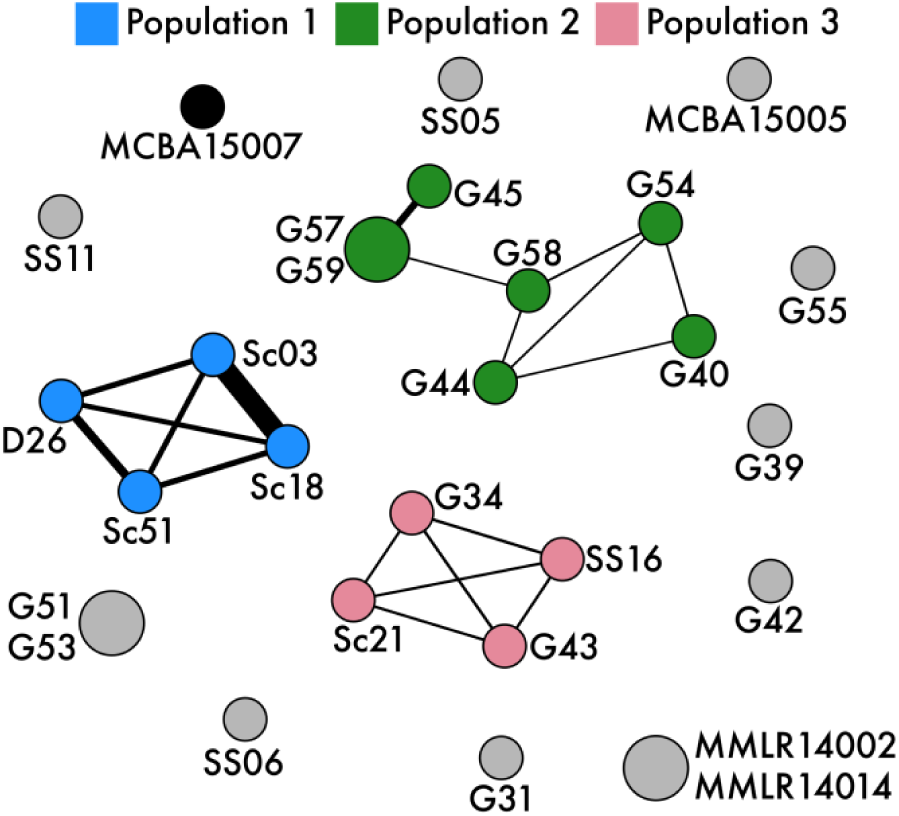
Recombination network across all pairwise strains. Thicker edges represent increased recombination between strains. Nodes are colored by population designation and node size indicates number of clonal clusters (strains too closely-related to differentiate recombination). D = Desert, Sc = Scrubland, G/MMLR = Grassland, SS = Salton Sea, MCBA = Boston

This approach enabled the identification of recombining populations that would otherwise be masked with traditional phylogenetic analyses. For example, two strains (MMLR14002/014) isolated from the grassland site five years prior share no recent recombination events (Fig. 2) despite sharing a high degree of phylogenetic relatedness and a common ancestral gene pool to strains within population 3 (Fig. 1). Additionally, the analysis revealed that the highly similar strains isolated across the continent from one another (from Boston, MA and a CA grassland; Fig. 1) were not connected by recent recombination events. Indeed, this conservative approach to estimate recombination events reduced most strains within the IB/C subclade to singleton nodes, suggesting that no recent recombination events connect these individuals to the three identified populations (Fig. 2), and that these strains are probably representatives of other, unsampled populations.

To confirm the effect of homologous recombination on the genetic diversity within subclade IB/C, we employed ClonalFrameML (Didelot and Wilson, 2015). Specifically, we concentrated on the r/m ratio at which nucleotides are substituted from either recombination or point mutations. Throughout the evolution of the IB/C subclade, recombination rates were generally low (r/m = 0.94), indicating barriers to gene flow and the occurrence of mutation accumulation within the subclade. However, when we assessed the rates of recombination within each population assignment, we found that homologous recombination rates to be high in populations 1 and 3 (r/m = 3.34 and 2.75, respectively) while population 2 (r/m = 1.62) had intermediate recombination rates (Supplementary Table 2). The observed r/m values are especially notable as terrestrial free-living bacteria have previously been shown to have low r/m values (r/m <1) (Vos and Didelot, 2008).

### Population Differentiation of the Flexible Genome

Based on the recombination networks, we expected that individuals within the same population would also share more flexible genes (genes not present in all strains) than individuals between different populations. The similarity between flexible gene content among strains was highly congruent with the population assignments (Fig. 3); strains within a population (ANOSIM; R = 0.88, p = 0.001) shared more flexible genes than expected by chance. We also observed that flexible gene content differed significantly by site (ANOSIM; R = 0.81, p < 0.01), suggesting that processes within and across locations are structuring the differences in the flexible genome within the subclade IB/C.

**FIG 3.**
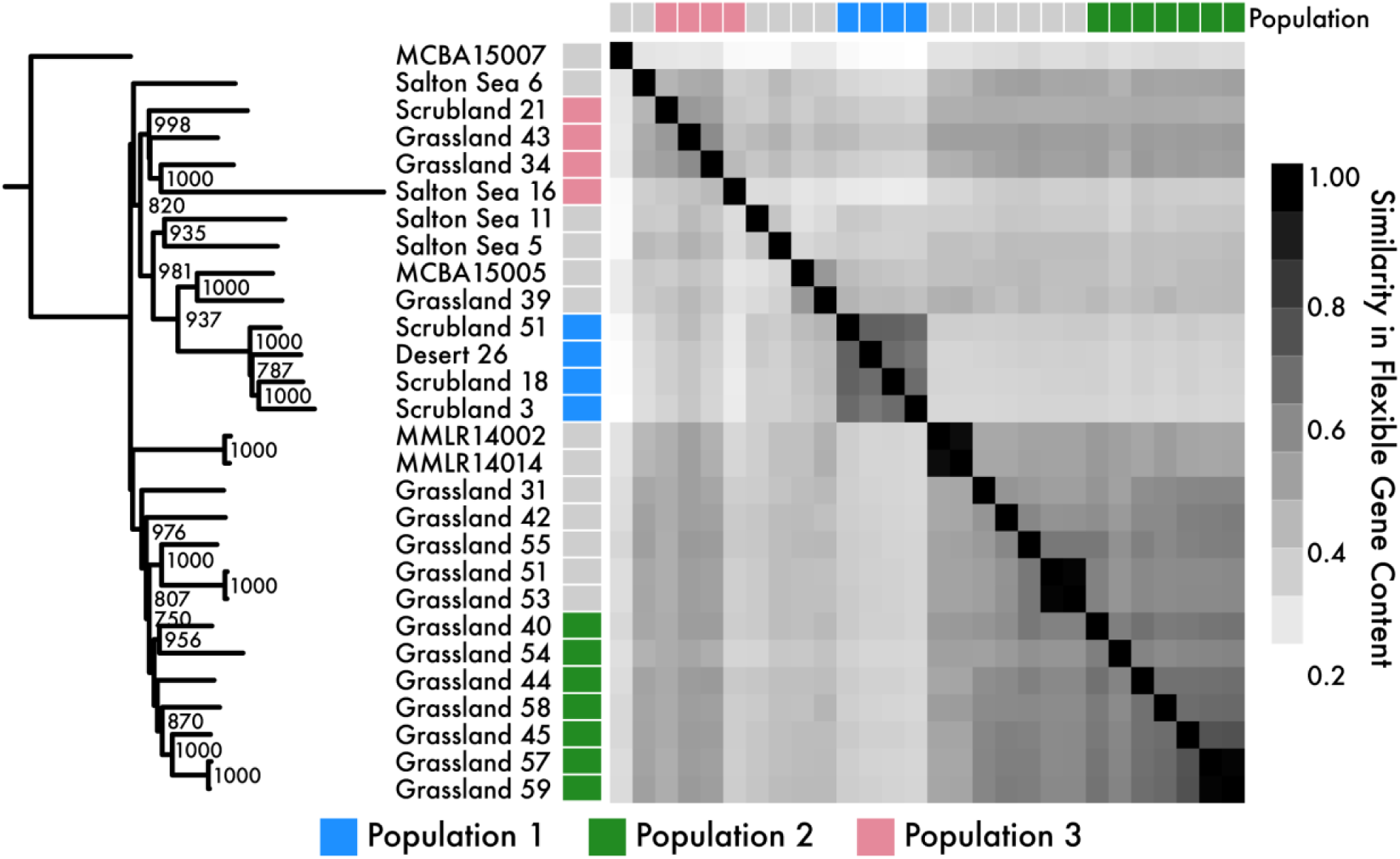
Flexible gene content similarity between strains. Tree is derived from a consensus neighbor-joining analysis showing only nodes with ≥750 support. Strains are colored by population assignments identified from the recombination network (Fig. 2).

The flexible genome also provides insights into the traits that distinguish populations. For example, flexible genes only present in all individuals within a particular population may have swept through the population by positive selection (Polz *et al.*, 2013). We searched for population-specific genes shared among all members and discovered that many were highly localized to a limited number of genomic regions. Specifically, 16 of 48 population-specific genes in population 1 were highly localized in the genome, while 4 of 6 population-specific genes in population 3 were localized (Fig. 4A). Additionally, these population-specific genes had reduced nucleotide diversity when compared to whole-genome measurements (Supplementary Figure S1), which can be indicative of relatively recent selective sweeps. These putative sweep regions may have arrived prior to population diversification and subsequently co-diversified, but, nonetheless, represent genomic regions harboring population-specific flexible genes. We did not detect any localization of population-specific genes in population 2, perhaps due to its lower rate of homologous recombination (Supplementary Table 2).

**FIG 4.**
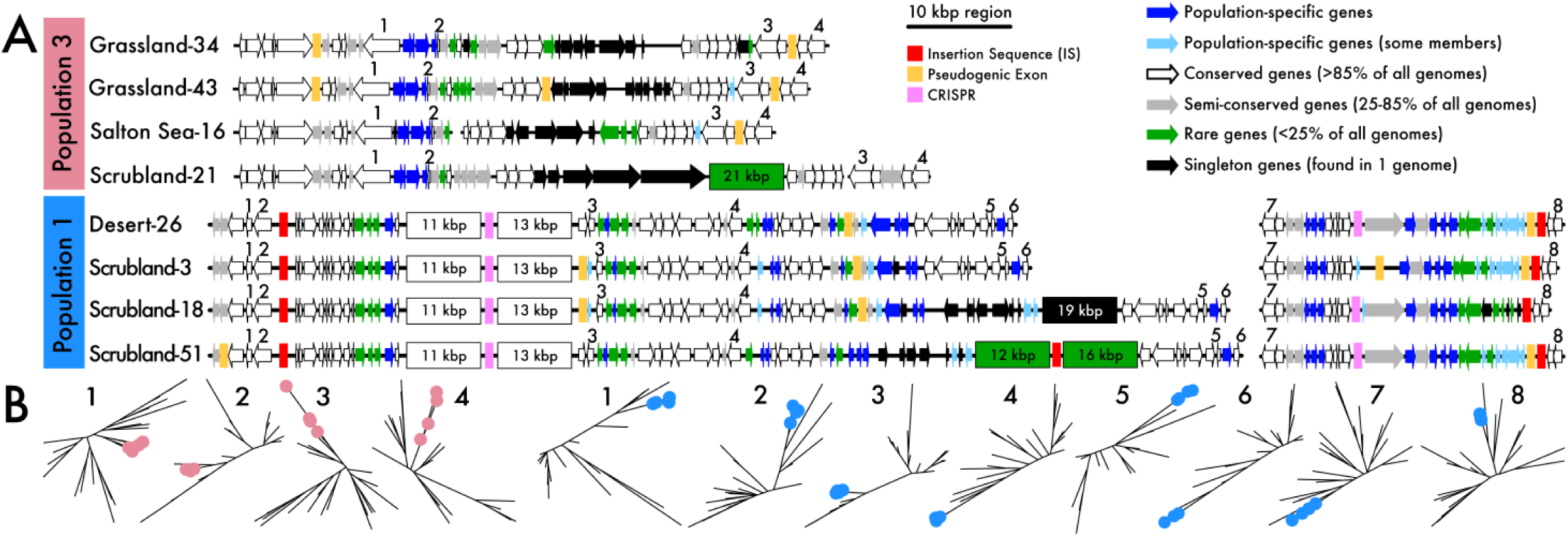
Highly structured genomic backbones across strains. (A) Population-specific genomic backbones within all individuals in populations 1 and 3. Population-specific genes (colored in blue) are consistently flanked by highly conserved regions (in white). Putative mobile elements are also designated in boxes along the chromosome. (B) Phylogenies of a subset of conserved genes (white arrows in panel A) flanking the population-specific regions colored by the strains in each respective population.

The flanking genomic regions surrounding the population-specific genes exhibited high genomic conservatism across all members in the population as well, suggesting these genomic regions may be hotspots for genetic exchange within the populations (Fig. 4A). While we did not detect phage or integrative and conjugative elements (ICEs), we did identify other mobile elements such as insertion sequences and clustered regularly interspaced short palindromic repeats (CRISPRs). Further, the regions were littered with pseudogenic exons, indicating the interruption of functional proteins due to recombining genomic segments. The genomic regions also contained rare (<25% of all members within Subclade IB/C) or strain-specific genes. In contrast to these variable regions, the flanking genes were highly conserved (shared by >85% of all members within Subclade IB/C) in nearly identical genetic architectures. Many of the conserved flanking core genes supported a strict monophyletic division of the population (Fig. 4B).

Most of the population-specific genes within the variable regions annotate as hypothetical proteins with some transcriptional regulators; however, other genes may be involved in differential use of environmental resources. For example, the regions contained a high number of metal uptake and transport proteins, along with glycoside hydrolase (GH) enzymes and glycosyltransferases that contribute to the breakdown of carbohydrates commonly found in leaf litter. To that end, we also observed a difference in the full genomic potential to degrade various carbohydrates in leaf litter between populations (ANOVA; p < 0.01). However, other predicted genomic traits (i.e. minimum generation time and optimal growth temperature) were indistinguishable between populations, most likely due to the calculation incorporating full genome-wide codon usage biases (Supplementary Figure S2).

## DISCUSSION

Our results suggest that both allopatric and sympatric processes are responsible for structuring populations of free-living soil bacteria across a regional climate gradient. This genetic resolution was possible by isolating a variety of *Curtobacterium* strains from the same habitat (leaf litter) across geographic locations (Chase *et al.*, 2018). Within the most abundant ecotype, Subclade IB/C, we quantified gene flow among closely-related, co-occurring lineages to identify distinct genetic populations of *Curtobacterium* across geographic distances. An analysis of the flexible genome confirmed that these populations are structured by gene flow discontinuities and provided additional evidence for population-specific adaptation. Finally, the distributional patterns of the populations suggest that both isolation-by-distance and isolation-by-environment contribute to *Curtobacterium* population structure. Thus, both dispersal limitation and local environmental adaptation contribute to the divergence among closely-related soil bacteria as observed in macroorganisms (Sexton *et al.*, 2014).

Previously, studies of two soil bacteria, *Streptomyces* and *Bradyrhizobium*, found continental-scale patterns consistent with allopatric diversification over distantly-related strains (<90% ANI) (VanInsberghe *et al.*, 2015; Andam *et al.*, 2016; Choudoir *et al.*, 2016). Further, clonal sympatric strains of the social bacterium *Myxococcus* were found to have barriers to recombination over cm distances in soil (Wielgoss *et al.*, 2016). By isolating strains within a single *Curtobacterium* ecological cluster at varying geographic scales, we could characterize the processes driving recent population divergence between both co-occurring strains and across regional spatial scales. As a comparison, we included two strains within this ecotype that were isolated from Boston, MA and found no recent recombination events connecting strains across continental scales (Fig. 2). Notably, along the regional climate gradient, we found that closely-related strains isolated from similar leaf litter communities were constrained in their geographic extent (mean geographic range of populations = 62.4 ± 100 km), suggesting that observed gene flow patterns is consistent with allopatric differentiation. However, we also observed multiple, genetically-distinct populations overlapping at three of the sites. Two of these populations were comprised of individuals from spatially distinct sites that remained connected by gene flow, suggesting isolation-by-distance is reduced at regional spatial scales. These results are contrary to previous work in fungal populations conducted at similar spatial scales; where fungal populations were highly structured by geography insomuch that genomic differences strongly reflected local site adaptations, a pattern consistent with strictly allopatric differentiation (Branco *et al.*, 2015; Amend *et al.*, 2010).

The presence of sympatric *Curtobacterium* populations can indicate the presence of an isolating mechanism to maintain the cohesiveness of co-occurring genetic lineages (Mayr, 2001). For instance, the efficiency of homologous recombination between bacteria can decrease exponentially with increasing sequence divergence (Fraser *et al.*, 2007). Alternatively, the presence of sympatric populations could signify that spatial barriers between the populations existed in the past but have since been removed without sufficient time for genetic homogenization. The flexible genome of *Curtobacterium* provides two lines of evidence for the former and, specifically, that the identified populations have remained genetically isolated due to ecological differentiation, as others have observed in bacterial populations (Shapiro and Polz, 2014). First, *Curtobacterium* populations shared more flexible genes within populations than between, suggesting that the populations represent cohesive, ecologically differentiated clusters (Fig. 3). Flexible genes are thought to contribute to differences in niche exploitation (Rodriguez-Valera and Ussery, 2012) and can contribute to small fitness differences among microhabitats (Cordero and Polz, 2014). For example, in the marine bacterium *Vibrio*, sympatric populations encoded habitat-specific genes (Shapiro *et al.*, 2012) between free-living and particle-associated populations (Yawata *et al.*, 2014). At a similar microscale, *Curtobacterium* populations may differentiate between leaf litter microhabitats caused by variability in resources such as metals and carbohydrate availability. Accordingly, we observed differences in carbohydrate degradation potential and observed population-specific genomic islands encoding genes related to physiological features.

The second line of evidence that sympatric populations are being maintained by ecological differences is that all individuals within populations shared highly conserved genomic backbones containing population-specific genes (Fig. 4). The population-specific genomic backbones consisted of both core genes exhibiting a strict monophyletic division and population-specific flexible genes indicating recent selective sweeps within a population. These patterns have been previously identified in marine bacterial populations of *Vibrio* (Shapiro *et al.*, 2012) and *Prochlorococcus* (Kashtan *et al.*, 2014) and the archaeon *Sulfolobus* (Cadillo-Quiroz *et al.*, 2012), where population-specific genomic regions were linked to small fitness differences and niche exploitation contributing to the coexistence of sympatric populations. Similarly, increased homologous recombination among strains of *Curtobacterium* populations could enable the rapid exchange of niche-adaptive genes for differential microhabitat specialization on leaf litter. This observation is consistent with isolation-by-environment where gene exchange rates among similar environments is higher than within geographic locations (Wang and Bradburd, 2014). Thus, the populations along the regional climate gradient seem to represent genetically-isolated lineages that are ecologically diverged by their partitioning microhabitats (within a location).

A major gap in our understanding of microbial diversity is the mechanisms contributing to the origin and maintenance of microbial diversification. Collectively, our results suggest a model for the recent microevolution of a soil bacterium. Similar to soil fungal populations and macroorganisms, free-living soil bacterial populations are geographically restricted. At the same time, distinct *Curtobacterium* populations may have also diverged to specialize on different leaf litter microhabitats, causing a reduction in gene flow between populations. Thus, overlapping populations are maintained within the same location, while also being connected via dispersal to individuals in other locations. Our results demonstrate that soil bacterial populations, similarly to those in other environments, are delineated by barriers to recombination where the proliferation of advantageous genes can spread in a population-specific manner (Whitaker *et al.*, 2005; Fraser *et al.*, 2007; Cadillo-Quiroz *et al.*, 2012; Shapiro *et al.*, 2012).

## MATERIALS AND METHODS

### Field Sites and *Curtobacterium* Strains

We downloaded 28 *Curtobacterium* genomes (Supplementary Table 1) from the National Center for Biotechnology Information (NCBI) [https://www.ncbi.nlm.nih.gov/] database that were previously isolated from leaf litter (Chase *et al.*, 2016), including a robust genomic dataset consisting of 26 strains from a climate gradient in southern California (Chase *et al.*, 2018). We included two additional strains within the same ecotype from outside Boston, MA to provide varying spatial scales for population comparisons. Protein-coding regions and gene annotations were derived from the NCBI prokaryotic genome annotation pipeline (Tatusova *et al.*, 2016). Genomes were screened for the presence of mobile elements by identifying integrating and conjugative elements (ICEs) with the ICEberg database (Bi *et al.*, 2011), prophage sequences using PhiSpy (Akhter *et al.*, 2012), insertion sequences (IS) with ISfinder (Siguier *et al.*, 2006), and CRISPR with CRISPRCasFinder (Couvin *et al.*, 2018).

### Evolutionary History of the Core Genome

We aligned all genomes using progressiveMauve (Darling *et al.*, 2010) to identify locally collinear blocks (LCBs) of genomic data. We identified 49,610 LCBs >1500 bp found across all 28 genomes that represented 1.28 Mbp of the core genome. This core genome alignment was used to perform a maximum likelihood bootstrap analysis using RAxML v8.2.10 (Stamatakis, 2014) under the general time reversal model with a gamma distribution for 100 replicates.

Using the core genome, we performed an initial analysis to infer the relative effects of recombination and mutation rates using ClonalFrameML v1.11 (Didelot and Wilson, 2015). Specifically, we attempted to reconstruct phylogenetic relationships by detecting regions of recombination across the phylogeny to provide an initial estimate for clonal genealogy.

Due to the weak clonal structure among strains, we sought to infer population structure from multilocus genotype data. First, we converted the core genome sequence data to a genotype matrix reflecting the distance between polymorphic sites of all individuals (https://github.com/xavierdidelot). We then used this genotype matrix to compute ancestry coefficients to delineate genetic clusters. Specifically, we employed sparse non-negative matrix factorization algorithms to estimate the cross-entropy parameter (Frichot *et al.*, 2014). Based on the cross-entropy criterion which best fit the statistical model, we designated the number of ancestral populations to K=4 to estimate individual admixture coefficients using the LEA package (Frichot and François, 2015) in the R software environment (Pinheiro *et al.*, 2011).

### Gene Flow and Recombination Networks

To differentiate between vertical transmission and recent recombination, we identified recent transfer events across all pairs of genomes using PopCOGenT (https://github.com/philarevalo/PopCOGenT). Briefly, we used a null model of sequence divergence to calculate the expected length distribution of identical genomic regions between strain pairs. Recently exchanged genes would enrich this distribution by introducing identical genomic regions that are longer and more frequent than expected. The extent of this enrichment is our measurement of recent transfer. Strains that were too closely related (<0.035% ANI divergence) to accurately assess recombination transfers were collapsed into clonal complexes. Finally, strains that were connected to any other strain in the recombination network were considered to be a part of the same recombining population. To confirm the importance of recombination events in structuring populations, we inferred the relative effects of recombination and mutations rates of the core genome (see above) within each population using ClonalFrameML.

### Population Genetic Analyses

To perform within population genetic analyses, we identified all orthologous protein-coding genes (orthologs) shared across all strains. Orthologs were initially predicted using ROARY (Page *et al.*, 2015) with a minimum sequence identity of 90% to ensure all possible orthologs were included across populations (Supplementary Figure 3A). The resulting 2193 orthologs shared across all strains were individually aligned with ClustalO v1.2.3 (Sievers *et al.*, 2011) and used to create a 2.14 Mbp concatenated nucleotide alignment. Note, the size of this alignment differs from the core genome alignment since genes do not necessarily need to be located on LCBs. To verify the effects of using a gene × gene approach on the core genome, we reconstructed the phylogenetic relationship of the concatenated alignment of all orthologous protein-coding genes, using RAxML v8.2.10 (Stamatakis, 2014) under the general time reversal model with a gamma distribution for 100 replicates, and compared to phylogeny derived from the Mauve core genome alignment (Supplementary Figure 3B). Next, all individual ortholog alignments were screened for complete codon reading frames (i.e. multiple of 3 bp) and the resulting 2137 genes were individually used to calculate nucleotide diversity within populations using the PopGenome package (Pfeifer *et al.*, 2014) in R, as outlined in (Lemieux *et al.*, 2016).

Predicted orthologs that were not shared across all strains represent the flexible genome (Supplementary Figure 3A). Using all identified orthologs, we computed a Jaccard distance between pairs of strains to estimate shared gene content. The distance matrix was used to generate a neighbor-joining tree based on 1000 re-samplings and to create a heatmap showing gene content similarity across strains. We tested the significance of gene content using an analysis of similarities (ANOSIM) for populations and site of isolation for 9999 permutations. In addition, we looked for orthologs that were unique to our populations. Specifically, we identified orthologs that were encoded by every member within a population and were not found in any member outside of the population. To reduce this list even further, we identified population-specific orthologs that were localized in genomic regions (<10 kbp separation).

### Analysis of Genomic Traits

We analyzed all genomic sequences for specific ecological traits that may contribute to population divergence. We concentrated on genomic traits related to growth strategies and substrate (i.e. carbohydrate) utilization that may be advantageous on leaf litter.

To infer growth strategies, we estimated minimum generation times (MGT) and optimal growth temperature (OGT). We predicted MGT by comparing codon-usage biases between highly expressed ribosomal proteins and all other encoded genes following a linear regression model (Vieira-Silva and Rocha, 2010)[equation 1].

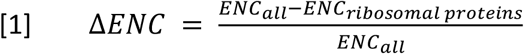

ENC = effective number of codons given %GC (Subramanian, 2008)

We analyzed each strain for the genomic potential to degrade various carbohydrates by searching the predicted coding-regions against the Pfam-A v30.0 database (Finn *et al.*, 2016) using HMMer (Finn *et al.*, 2011). Identified protein families were reduced to only known protein families that encode for glycoside hydrolase (GH) and carbohydrate binding module (CBM) proteins as described in (Chase *et al.*, 2016).

## ACKNOWLEDGMENTS

The authors thank Claudia Weihe, Chamee Moua, and Michaeline Albright for their assistance in sample collection and strain isolation. We also thank Brandon Gaut for invaluable insight into data analysis and interpretation, Sarai Finks, Kendra Walters, Cynthia Rodriguez, and the rest of the Martiny Lab for helpful comments, and Xavier Didelot and Kevin Bonham for software assistance. This work was supported by an US Department of Education Graduate Assistance of National Need (GAANN) fellowship to ABC, the US Department of Energy, Office of Science Graduate Student Research (SCGSR) fellowship to ABC, and the US Department of Energy, Office of Science, Office of Biological and Environmental Research (DE-SC0016410) to JBHM.

## Notes

#### Summary of Updates

Clarified the discussion; revised abstract

## REFERENCES

Akhter S, Aziz RK, Edwards RA. (2012). PhiSpy: a novel algorithm for finding prophages in bacterial genomes that combines similarity-and composition-based strategies. Nucleic Acids Res 40: e126–e126.

Amend A, Garbelotto M, Fang Z, Keeley S. (2010). Isolation by landscape in populations of a prized edible mushroom Tricholoma matsutake. Conserv Genet 11: 795–802.

Andam CP, Doroghazi JR, Campbell AN, Kelly PJ, Choudoir MJ, Buckley DH. (2016). A latitudinal diversity gradient in terrestrial bacteria of the genus Streptomyces. MBio 7: e02200–15.

Arevalo P, VanInsberghe D, Elsherbini J, Gore J, Polz MF. (2019). A reverse ecology approach based on a biological definition of microbial populations. Cell.

Bi D, Xu Z, Harrison EM, Tai C, Wei Y, He X, et al. (2011). ICEberg: a web-based resource for integrative and conjugative elements found in Bacteria. Nucleic Acids Res 40: D621–D626.

Branco S, Gladieux P, Ellison CE, Kuo A, LaButti K, Lipzen A, et al. (2015). Genetic isolation between two recently diverged populations of a symbiotic fungus. Mol Ecol 24: 2747–2758.

Cadillo-Quiroz H, Didelot X, Held NL, Herrera A, Darling A, Reno ML, et al. (2012). Patterns of gene flow define species of thermophilic Archaea. PLoS Biol 10: e1001265.

Chase AB, Arevalo P, Polz MF, Berlemont R, Martiny JBH. (2016). Evidence for ecological flexibility in the cosmopolitan genus Curtobacterium. Front Microbiol 7: 1874.

Chase AB, Gomez-Lunar Z, Lopez AE, Li J, Allison SD, Martiny AC, et al. (2018). Emergence of soil bacterial ecotypes along a climate gradient. Environ Microbiol 20: 4112–4126.

Chase AB, Karaoz U, Brodie EL, Gomez-Lunar Z, Martiny AC, Martiny JBH. (2017). Microdiversity of an Abundant Terrestrial Bacterium Encompasses Extensive Variation in Ecologically Relevant Traits. MBio 8: e01809–17.

Chase AB, Martiny JBH. (2018). The importance of resolving biogeographic patterns of microbial microdiversity. Microbiol Aust 39: 5–8.

Choudoir MJ, Buckley DH. (2018). Phylogenetic conservatism of thermal traits explains dispersal limitation and genomic differentiation of Streptomyces sister-taxa. ISME J 1.

Choudoir MJ, Doroghazi JR, Buckley DH. (2016). Latitude delineates patterns of biogeography in terrestrial Streptomyces. Environ Microbiol 18: 4931–4945.

Cohan FM. (2001). Bacterial species and speciation. Syst Biol 50: 513–524.

Cordero OX, Polz MF. (2014). Explaining microbial genomic diversity in light of evolutionary ecology. Nat Rev Microbiol 12: 263–273.

Couvin D, Bernheim A, Toffano-Nioche C, Touchon M, Michalik J, Néron B, et al. (2018). CRISPRCasFinder, an update of CRISRFinder, includes a portable version, enhanced performance and integrates search for Cas proteins. Nucleic Acids Res.

Cui Y, Yang X, Didelot X, Guo C, Li D, Yan Y, et al. (2015). Epidemic clones, oceanic gene pools, and eco-LD in the free living marine pathogen Vibrio parahaemolyticus. Mol Biol Evol 32: 1396–1410.

Darling AE, Mau B, Perna NT. (2010). progressiveMauve: multiple genome alignment with gene gain, loss and rearrangement. PLoS One 5: e11147.

Didelot X, Wilson DJ. (2015). ClonalFrameML: efficient inference of recombination in whole bacterial genomes. PLoS Comput Biol 11: e1004041.

van Elsas JD, Bailey MJ. (2002). The ecology of transfer of mobile genetic elements. FEMS Microbiol Ecol 42: 187–197.

Finn RD, Clements J, Eddy SR. (2011). HMMER web server: Interactive sequence similarity searching. Nucleic Acids Res 39: 29–37.

Finn RD, Coggill P, Eberhardt RY, Eddy SR, Mistry J, Mitchell AL, et al. (2016). The Pfam protein families database: Towards a more sustainable future. Nucleic Acids Res 44: D279–D285.

Fraser C, Hanage WP, Spratt BG. (2007). Recombination and the nature of bacterial speciation. Science (80-) 315: 476–480.

Frichot E, François O. (2015). LEA: an R package for landscape and ecological association studies. Methods Ecol Evol 6: 925–929.

Frichot E, Mathieu F, Trouillon T, Bouchard G, François O. (2014). Fast and efficient estimation of individual ancestry coefficients. Genetics genetics-113.

Hunt DE, David LA, Gevers D, Preheim SP, Alm EJ, Polz MF. (2008). Resource partitioning and sympatric differentiation among closely related bacterioplankton. Science (80-) 320: 1081–1085.

Jain C, Rodriguez-R LM, Phillippy AM, Konstantinidis KT, Aluru S. (2017). High-throughput ANI analysis of 90K prokaryotic genomes reveals clear species boundaries. bioRxiv 225342.

Johnson ZI, Zinser ER, Coe A, McNulty NP, Woodward EMS, Chisholm SW. (2006). Niche partitioning among Prochlorococcus ecotypes along ocean-scale environmental gradients. Science (80-) 311: 1737–1740.

Kashtan N, Roggensack SE, Rodrigue S, Thompson JW, Biller SJ, Coe A, et al. (2014). Single-cell genomics reveals hundreds of coexisting subpopulations in wild Prochlorococcus. Science (80-) 344: 416–420.

Lemieux JE, Tran AD, Freimark L, Schaffner SF, Goethert H, Andersen KG, et al. (2016). A global map of genetic diversity in Babesia microti reveals strong population structure and identifies variants associated with clinical relapse. Nat Microbiol 1: 16079.

Mayr E. (2001). What evolution is. Science Masters Series.

Nannipieri P, Ascher J, Ceccherini M, Landi L, Pietramellara G, Renella G. (2003). Microbial diversity and soil functions. Eur J Soil Sci 54: 655–670.

Page AJ, Cummins CA, Hunt M, Wong VK, Reuter S, Holden MTG, et al. (2015). Roary: rapid large-scale prokaryote pan genome analysis. Bioinformatics 31: 3691–3693.

Pfeifer B, Wittelsbürger U, Ramos-Onsins SE, Lercher MJ. (2014). PopGenome: an efficient Swiss army knife for population genomic analyses in R. Mol Biol Evol 31: 1929–1936.

Pinheiro J, Bates D, DebRoy S, Sarkar D. (2011). R Development Core Team. 2010. nlme: linear and nonlinear mixed effects models. R package version 3. 1–97. R Found Stat Comput Vienna.

Polz MF, Alm EJ, Hanage WP. (2013). Horizontal gene transfer and the evolution of bacterial and archaeal population structure. Trends Genet 29: 170–175.

Ranjard L, Richaume A. (2001). Quantitative and qualitative microscale distribution of bacteria in soil. Res Microbiol 152: 707–716.

Ravenhall M, Škunca N, Lassalle F, Dessimoz C. (2015). Inferring horizontal gene transfer. PLoS Comput Biol 11: e1004095.

Rocha EPC. (2018). Neutral theory, microbial practice: challenges in bacterial population genetics. Mol Biol Evol 35: 1338–1347.

Rodriguez-R LM, Konstantinidis KT. (2014). Bypassing cultivation to identify bacterial species. Microbe 9: 111–118.

Rodriguez-Valera F, Ussery DW. (2012). Is the pan-genome also a pan-selectome? F1000Research 1.

Sexton JP, Hangartner SB, Hoffmann AA. (2014). Genetic isolation by environment or distance: which pattern of gene flow is most common? Evolution (N Y) 68: 1–15.

Shapiro BJ, Friedman J, Cordero OX, Preheim SP, Timberlake SC, Szabó G, et al. (2012). Population genomics of early events in the ecological differentiation of bacteria. Science (80-) 336: 48–51.

Shapiro BJ, Leducq J-B, Mallet J. (2016). What is speciation? PLoS Genet 12: e1005860.

Shapiro BJ, Polz MF. (2014). Ordering microbial diversity into ecologically and genetically cohesive units. Trends Microbiol 22: 235–247.

Sheppard SK, McCarthy ND, Falush D, Maiden MCJ. (2008). Convergence of Campylobacter species: implications for bacterial evolution. Science (80-) 320: 237–239.

Sievers F, Wilm A, Dineen D, Gibson TJ, Karplus K, Li W, et al. (2011). Fast, scalable generation of high-quality protein multiple sequence alignments using Clustal Omega. Mol Syst Biol 7: 539.

Siguier P, Pérochon J, Lestrade L, Mahillon J, Chandler M. (2006). ISfinder: the reference centre for bacterial insertion sequences. Nucleic Acids Res 34: D32–D36.

Stamatakis A. (2014). RAxML version 8: A tool for phylogenetic analysis and post-analysis of large phylogenies. Bioinformatics 30: 1312–1313.

Subramanian S. (2008). Nearly neutrality and the evolution of codon usage bias in eukaryotic genomes. Genetics 178: 2429–2432.

Tatusova T, DiCuccio M, Badretdin A, Chetvernin V, Nawrocki EP, Zaslavsky L, et al. (2016). NCBI prokaryotic genome annotation pipeline. Nucleic Acids Res 44: 6614–6624.

VanInsberghe D, Maas KR, Cardenas E, Strachan CR, Hallam SJ, Mohn WW. (2015). Non-symbiotic Bradyrhizobium ecotypes dominate North American forest soils. ISME J 9: 2435.

Vieira-Silva S, Rocha EPC. (2010). The systemic imprint of growth and its uses in ecological (meta) genomics. PLoS Genet 6: e1000808.

Vos M, Didelot X. (2008). A comparison of homologous recombination rates in bacteria and archaea. Isme J 3: 199.

Wang IJ, Bradburd GS. (2014). Isolation by environment. Mol Ecol 23: 5649–5662.

Ward DM, Cohan FM, Bhaya D, Heidelberg JF, Kühl M, Grossman A. (2008). Genomics, environmental genomics and the issue of microbial species. Heredity (Edinb) 100: 207.

Whitaker RJ, Grogan DW, Taylor JW. (2003). Geographic barriers isolate endemic populations of hyperthermophilic archaea. Science (80-) 301: 976–978.

Whitaker RJ, Grogan DW, Taylor JW. (2005). Recombination shapes the natural population structure of the hyperthermophilic archaeon Sulfolobus islandicus. Mol Biol Evol 22: 2354–2361.

Wielgoss S, Didelot X, Chaudhuri RR, Liu X, Weedall GD, Velicer GJ, et al. (2016). A barrier to homologous recombination between sympatric strains of the cooperative soil bacterium Myxococcus xanthus. ISME J.

Wright S. (1943). Isolation by distance. Genetics 28: 114.

Yawata Y, Cordero OX, Menolascina F, Hehemann J-H, Polz MF, Stocker R. (2014). Competition–dispersal tradeoff ecologically differentiates recently speciated marine bacterioplankton populations. Proc Natl Acad Sci 111: 5622–5627.

Zwirglmaier K, Jardillier L, Ostrowski M, Mazard S, Garczarek L, Vaulot D, et al. (2008). Global phylogeography of marine Synechococcus and Prochlorococcus reveals a distinct partitioning of lineages among oceanic biomes. Environ Microbiol 10: 147–161.

