## Supplemental Figures and Tables for "Sympatric and allopatric differentiation delineates population structure in free-living terrestrial bacteria"

### SUPPLEMENTAL MATERIAL

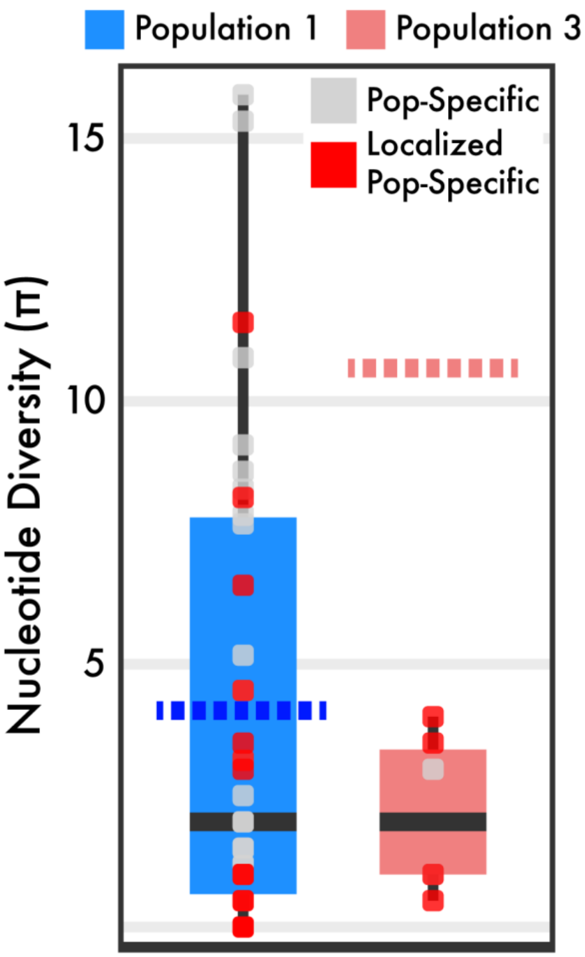


**FIG S1.** Boxplots show nucleotide diversity (π) across all population-specific genes (present in all members within the population). Each point is a population-specific gene and is colored whether the gene is localized in the genomic region displayed in Fig. 4. The dashed line shows the genome-wide average ($\pi_{MEAN}$) of each strain in the population across all core genes within subclade IB/C.

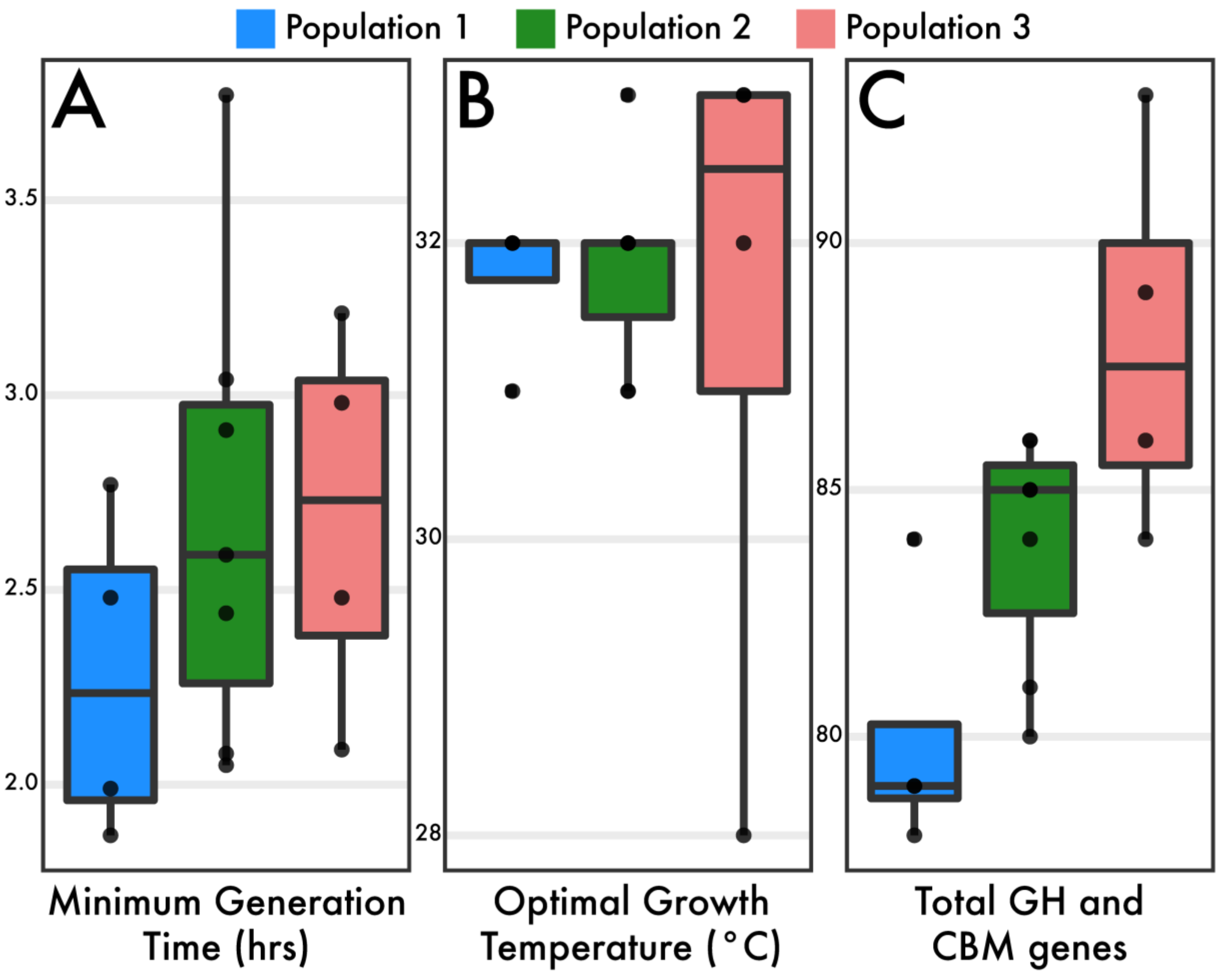


**FIG S2.** Distributions of predicted genomic traits in strains belonging to populations. Traits include: **(A)** minimum generation time (hrs), **(B)** optimal growth temperature (°C), and **(C)** total abundance of glycoside hydrolase (GH) and carbohydrate binding module (CBM) proteins.

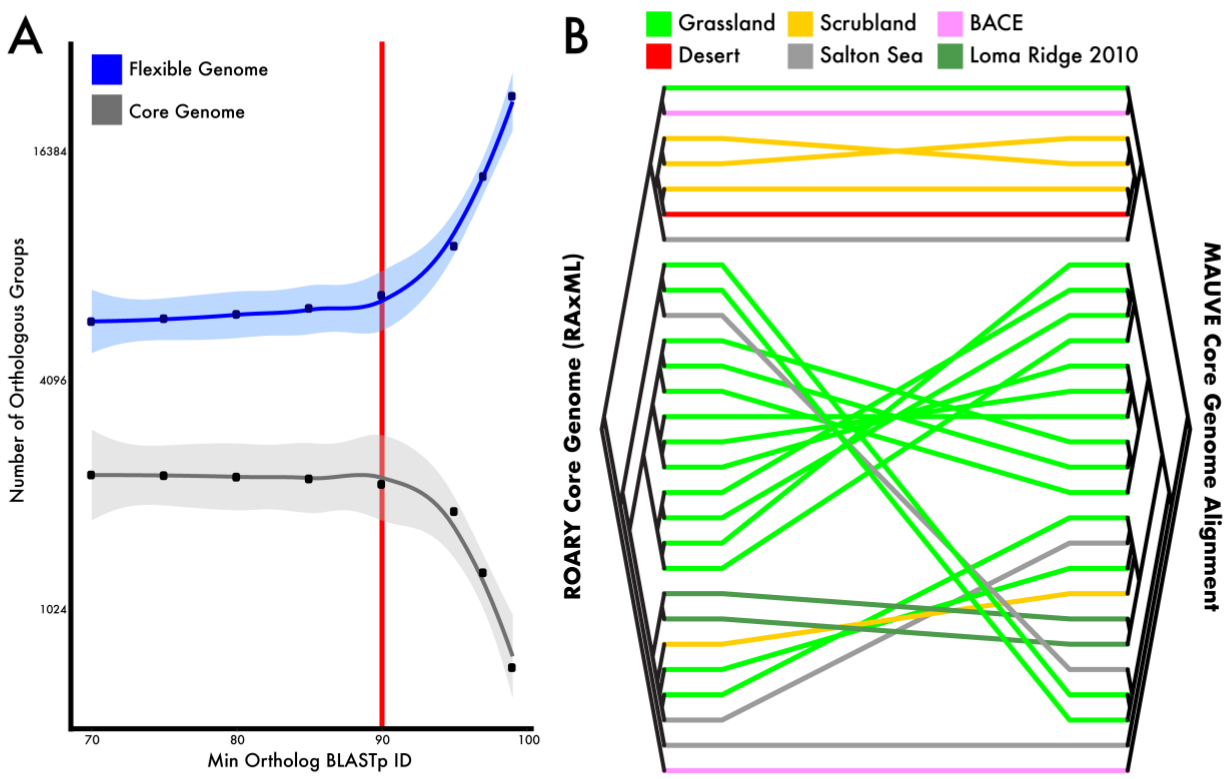


**FIG S3.** Breakdown of orthologous protein groups derived from all strains. **(A)** Number of identified orthologous protein groups in both the core and flexible genome based on initial clustering of proteins. **(B)** Cladogram comparison of core genes (N=2193 orthologous proteins) and core genome alignment (defined as locally collinear blocks). Terminal branches are colored by geographic location with lines connecting identical strains in each respective cladogram.

**TABLE S1.** Genomic and geographic characteristics of strains.

| Genome Name | BioSample ID | Location | Lat-Long | Isolation Year | Genome Length (bp) | %GC |
|---|---|---|---|---|---|---|
| MCBA15005 | SAMN05736482 | Boston, MA, USA | 42.38, -71.21 | 2015 | 3772244 | 69 |
| MCBA15007 | SAMN05736484 | Boston, MA, USA | 42.38, -71.21 | 2015 | 3768639 | 69 |
| MMLR14002 | SAMN05736479 | Loma Ridge, CA, USA | 33.74, -117.69 | 2011 | 3808678 | 70 |
| MMLR14014 | SAMN05736491 | Loma Ridge, CA, USA | 33.74, -117.69 | 2011 | 3822836 | 69 |
| Desert-26 | SAMN09009029 | Deep Canyon, CA, USA | 33.65, -116.37 | 2016 | 3645388 | 70 |
| Grassland-31 | SAMN09009040 | Loma Ridge, CA, USA | 33.74, -117.69 | 2016 | 3777956 | 71 |
| Grassland-34 | SAMN09009042 | Loma Ridge, CA, USA | 33.74, -117.69 | 2016 | 3820963 | 70 |
| Grassland-39 | SAMN09009044 | Loma Ridge, CA, USA | 33.74, -117.69 | 2016 | 3819069 | 70 |
| Grassland-40 | SAMN09009045 | Loma Ridge, CA, USA | 33.74, -117.69 | 2016 | 3761440 | 71 |
| Grassland-42 | SAMN09009046 | Loma Ridge, CA, USA | 33.74, -117.69 | 2016 | 3838097 | 71 |
| Grassland-43 | SAMN09009047 | Loma Ridge, CA, USA | 33.74, -117.69 | 2016 | 3695116 | 71 |
| Grassland-44 | SAMN09009048 | Loma Ridge, CA, USA | 33.74, -117.69 | 2016 | 3789494 | 71 |
| Grassland-45 | SAMN09009049 | Loma Ridge, CA, USA | 33.74, -117.69 | 2016 | 3764851 | 70 |
| Grassland-51 | SAMN09009050 | Loma Ridge, CA, USA | 33.74, -117.69 | 2016 | 3785722 | 71 |
| Grassland-53 | SAMN09009051 | Loma Ridge, CA, USA | 33.74, -117.69 | 2016 | 3786104 | 71 |
| Grassland-54 | SAMN09009052 | Loma Ridge, CA, USA | 33.74, -117.69 | 2016 | 3927711 | 70 |
| Grassland-55 | SAMN09009053 | Loma Ridge, CA, USA | 33.74, -117.69 | 2016 | 3752819 | 71 |
| Grassland-57 | SAMN09009054 | Loma Ridge, CA, USA | 33.74, -117.69 | 2016 | 3728355 | 71 |
| Grassland-58 | SAMN09009055 | Loma Ridge, CA, USA | 33.74, -117.69 | 2016 | 3800007 | 71 |
| Grassland-59 | SAMN09009056 | Loma Ridge, CA, USA | 33.74, -117.69 | 2016 | 3729778 | 71 |
| Salton Sea-11 | SAMN09009061 | Salton Sea, CA, USA | 33.33, -115.84 | 2016 | 3632003 | 70 |
| Salton Sea-16 | SAMN09009063 | Salton Sea, CA, USA | 33.33, -115.84 | 2016 | 4357272 | 70 |
| Salton Sea-5 | SAMN09009064 | Salton Sea, CA, USA | 33.33, -115.84 | 2016 | 3744522 | 70 |
| Salton Sea-6 | SAMN09009065 | Salton Sea, CA, USA | 33.33, -115.84 | 2016 | 3739706 | 70 |
| Scrubland-18 | SAMN09009071 | Pinon Flats, CA, USA | 33.61, -116.46 | 2016 | 3658022 | 70 |
| Scrubland-21 | SAMN09009073 | Pinon Flats, CA, USA | 33.61, -116.46 | 2016 | 3833068 | 70 |
| Scrubland-3 | SAMN09009074 | Pinon Flats, CA, USA | 33.61, -116.46 | 2016 | 3695248 | 70 |
| Scrubland-51 | SAMN09009079 | Pinon Flats, CA, USA | 33.61, -116.46 | 2016 | 3558624 | 71 |

**TABLE S2.** Ratio of nucleotide substitutions from recombination to point mutations (r/m).

| | | Population 1 | Population 2 | Population 3 | Subclade IB/C |
|---|---|---|---|---|---|
| N | Number of individuals in population | 4 | 7 | 4 | 28 |
| A | Geographic range of population (km) | 0 - 9.46 | 0 | 0 - 177.75 | 0 - 4032 |
| R/θ | ratio of recombination and mutation rates | 0.80 | 0.66 | 0.60 | 0.25 |
| δ | mean length of recombined fragments (bp) | 323.93 | 76.57 | 146.41 | 65.41 |
| ν | average divergence between donor/recipient | 0.01 | 0.03 | 0.03 | 0.06 |
| r/m | Relative effect of recombination and mutation | 3.34 | 1.62 | 2.75 | 0.94 |
